# A primary human muscle cell-based assay for detecting myasthenia gravis autoantibody binding and assessing AChR cluster impairment

**DOI:** 10.64898/2026.08.10.743478

**Authors:** Marlene Wolfsgruber, Anna-Sophie Zimmermann, Katja Starnberger, Tereza Duckova, Omar Keritam, Alexander Wöhrleitner, Rosa Weng, Paolo Doksani, Marta Rocha, Nina Matus, Katharina Tripkovic, Minal Pervez, Paloma Fernandes-Rosenegger, Francisca Faber, Cansu Elmas, Miriam Fichtner, Michelangelo Maestri Tassoni, Hakan Cetin, Romana Höftberger, Fritz Zimprich, Ruth Herbst, Christian Albrecht, Sarah Hoffmann, Lukas Weigl, Lilli Winter, Inga Koneczny

## Abstract

Myasthenia gravis (MG) is an autoimmune disease caused by pathogenic autoantibodies against proteins at the neuromuscular junction (NMJ). The diagnosis and clinical management of MG patients largely relies on the detection of antigen-specific autoantibodies targeting acetylcholine receptor (AChR) or muscle-specific kinase (MuSK). Yet a subset of patients remains seronegative for known MG autoantibodies, highlighting a critical need for alternative approaches to identify pathogenic NMJ antibodies. We established a new human *in vitro* model of the NMJ based on primary human muscle cells that recapitulates key features of the NMJ: differentiation to myotubes, expression of key NMJ proteins and formation of postsynaptic AChR clusters in response to agrin stimulation. The model allows new insights into myogenesis and genetic muscle diseases, and the new muscle cell-based assay (CBA) detected autoantibodies in sera from patients with AChR- and MuSK-positive MG with 96.43% sensitivity and 100% specificity, while healthy control sera showed no reactivity. Incubation with patient sera significantly reduced AChR clustering compared to controls, demonstrating functional pathogenic effects. Thus, we established a physiologically relevant human NMJ model that enables detection and functional characterization of neuromuscular autoantibodies. This novel approach addresses a key limitation of current antigen-specific diagnostics and provides a method for improved detection and characterization of MG antibodies, independent of antigen specificity.

**One Sentence Summary:** We established a postsynaptic human *in vitro* neuromuscular junction model to assess binding and pathogenicity of MG autoantibodies.

**Key messages:** *What is already known on this topic?:* Current diagnosis of myasthenia gravis (MG) relies largely on the detection of antigen-specific autoantibodies against AChR and MuSK, leaving a clinically relevant subset of patients seronegative.

*What are the new findings?:* We established a physiologically relevant human in vitro neuromuscular junction model based on primary human muscle cells and developed a novel muscle cell-based assay (CBA) for the detection of neuromuscular autoantibodies.

*How might this impact on clinical practice or future developments?:* The CBA detected autoantibodies in patients with AChR- or MuSK-positive MG with high sensitivity and specificity and demonstrated their functional pathogenic effects on AChR clustering. This antigen-independent approach may improve the detection and functional characterization of MG autoantibodies, particularly in patients who are seronegative in current diagnostic assays. 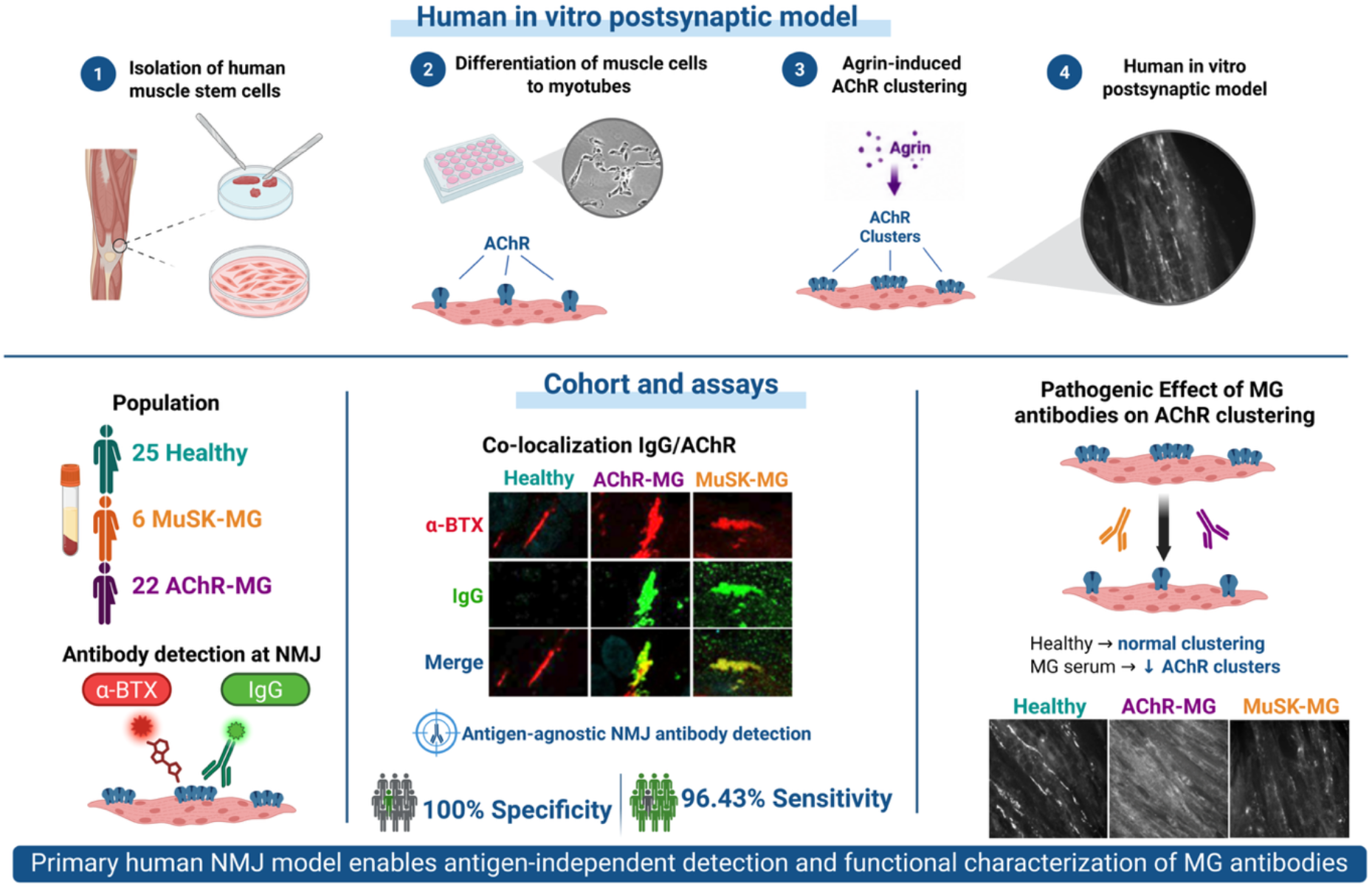

## INTRODUCTION

Myasthenia gravis (MG) is an autoimmune disease of the muscle, hallmarked by severe fatigable skeletal muscle weakness and caused by pathogenic autoantibodies that bind to proteins at the neuromuscular junction (NMJ) (1, 2). The NMJ is essential for the function of skeletal muscle, as motoneurons transmit the signal for muscle contraction via the neurotransmitter acetylcholine (ACh), which binds to postsynaptic acetylcholine receptors (AChR) in the muscle membrane (3). During NMJ development, neural agrin (4, 5) binds to its co-receptor low-density lipoprotein receptor-related protein 4 (LRP4) at the postsynaptic membrane, which in turn binds and activates muscle-specific kinase (MuSK), the master regulator of neuromuscular development (6–9). Binding of the adaptor protein DOK7 leads to the full activation of MuSK (10–12), and downstream signaling induces AChR clustering (13, 14) via the scaffolding protein rapsyn (15). In approximately 85% of the patients, pathogenic MG antibodies target the AChR, inducing complement activation, antigenic modulation or blocking (16–20). In about 5% of patients, antibodies against MuSK lead to a block of LRP4-MuSK interaction and reduced AChR clustering (21, 22). Both cases effectively lead to a reduction of functional AChRs and impaired neuromuscular transmission. Further antigens have been proposed as targets of autoantibodies in MG patients, including LRP4, collagen Q (ColQ) and agrin (23–28). However, a substantial number of patients (5-15%) remain seronegative (SNMG) for MG antibodies (29–32). The detection of autoantibodies in MG is essential for the diagnosis and guidance of treatment decisions, and the lack of known antigenic targets in SNMG is a challenge for the clinical management of the patients, warranting new approaches for antigen-agnostic antibody detection. Further, seropositive MG is a heterogenous disease, including different NMJ antigens and epitopes, different IgG subclasses, antibody valencies and pathogenic effector mechanisms at the NMJ (33), which are incompletely understood and require further characterization. To date, studies on functional effects of MG antibodies rely mostly on the use of C2C12 mouse muscle cells (34, 35), but rodent cells lack physiological relevance for the study of human disease (36, 37). Human rhabdomyosarcoma cell lines such as TE671 or CN21 (*38–42*) are of limited use as well, as they express tumor-associated mutations and lack differentiation potential. Immortalization and long-term *in vitro* culture can induce genomic alterations, shifts in gene expression profiles, and functional changes (*43–45*), making cell lines a less accurate compared with primary cells. We thus developed a new human *in vitro* model of the NMJ based on primary human muscle cells, with a cost-efficient isolation and culture protocol, a high yield of muscle cells that could be cryopreserved, differentiated to myotubes and robustly formed agrin-induced AChR clusters *in vitro*. Further, we could demonstrate that the model was suitable to detect MG autoantibodies with high specificity and sensitivity and was useful to study pathogenic effects of MG antibodies on AChR clustering. Beyond the field of MG, the model enables the study of genetic diseases of the muscle and NMJ, as it allows the characterization of patient-specific muscle cells.

## RESULTS

### Isolation of primary human muscle cells

We obtained human muscle as surgical waste from anterior cruciate ligament (ACL) reconstruction surgery from six healthy adult donors (D1-D6, two female, four males, median age: 32.5 years (range: 20-56), **Table S1**). The muscle stem cells were isolated as described previously (*46, 47*), based on the isolation procedure by Brinkmeier (*48*), within 24 hours of surgery. Non-muscle tissue was removed and muscle was dissociated by mechanical and enzymatic procedures (**Fig. 1A**). The cell slurry was cultured in growth medium established by Baroffio (*49*). As expected, during the first days, degradation of cellular debris was observed (**Fig. 1B**), thus the plates were kept in quarantine boxes to separate the cells from other cell lines in the incubator and prevent contamination. After 3-4 days, the first attaching cells could be observed on the plate (**Fig. 1B**), on day 6 the medium was changed. The cell colonies were detached when reaching a local confluence of 40% and further cultured at densities of 800–3500 cells/cm². P0 was defined when the cells were dense enough to be plated in a 100mm cell culture plate. The cells had a doubling time of 24 hours and were passaged at 40% confluence, allowing for a yield of around 4x10⁵ cells per 100mm plate.

**Fig. 1:**
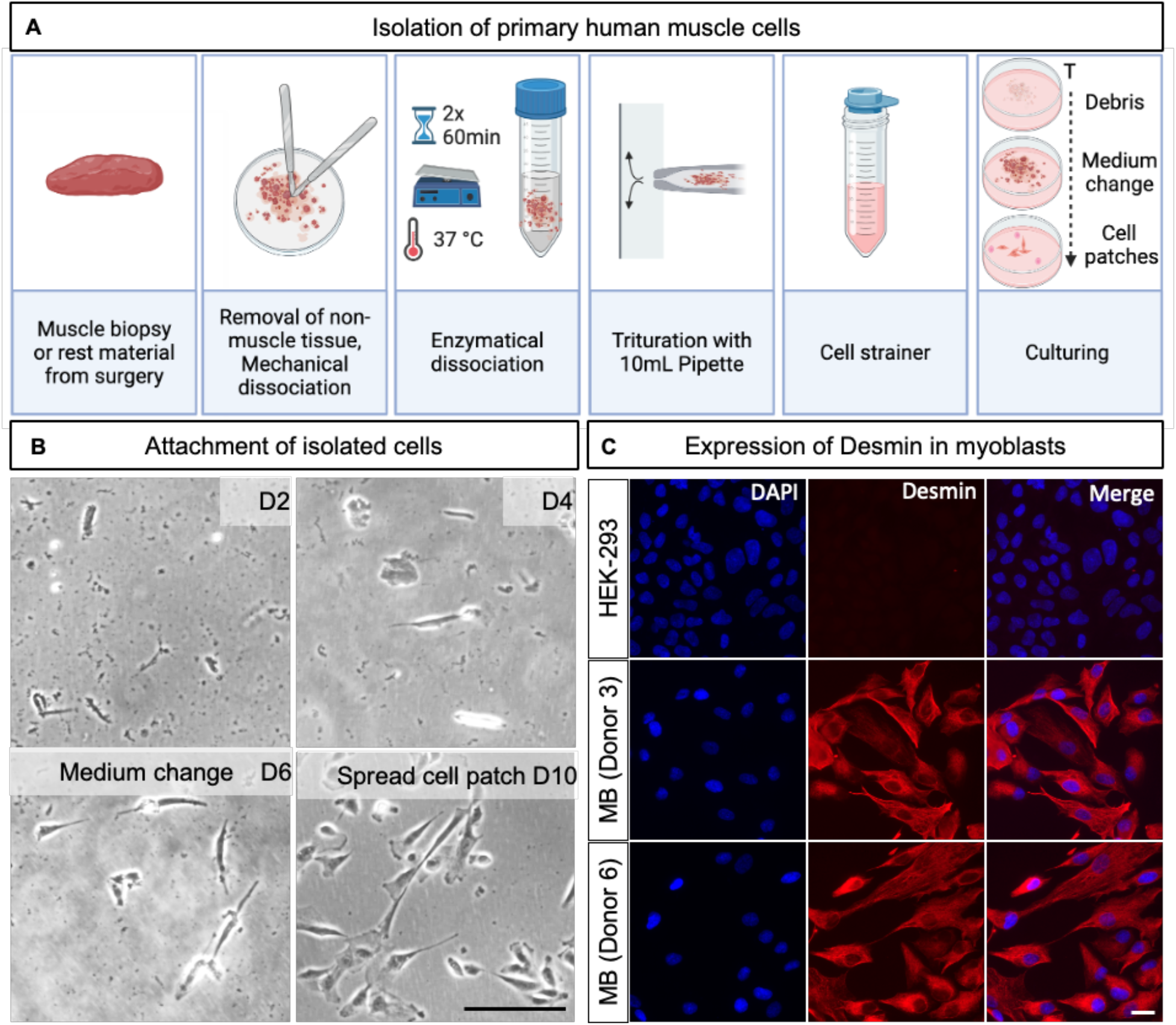
Isolation of primary human muscle cells. **(A)** Isolation strategy. **(B)** Brightfield microscopy images of culture plate post isolation day 2-10 (D2-D10). D2: Only cellular debris was visible in the initial days. D4: The first adherent cells were observed. D6: Medium change. D10: The cells reached approximately 40% confluency at day 10-12 post isolation. The scale bar indicates 100 µm. **(C)** Immunocytochemical staining of the intermediate filament protein desmin (visualized in red by AF594-conjugated secondary antibody) validated myoblast identity in isolated human cells (shown D3 and D6), but not HEK293 cells (negative control). DAPI staining was used to visualize the cell nuclei in blue. N = 3 for all 6 donors; the scale bar indicates 20 µm.

The cell population could be rapidly expanded, as each plate yielded enough cells to seed four new cell culture dishes, leading to approximately 64 vials by passage three, which could be frozen and stored in liquid nitrogen. After thawing, the cells were passaged once before use in experimental procedures. The primary human cells, but not HEK293 cells (negative control), were positive for the expression of the muscle-specific intermediate filament protein desmin, validating muscle cell identity (**Fig. 1C**).

### Evaluation of muscle cell differentiation and AChR clustering potential

Incubation of myoblasts with differentiation medium induced cell fusion, and the cells differentiated to myotubes within five to six days, with multiple nuclei per myotube visible in bright field microscopy (**Fig. 2A**, Day 6, arrowheads showing multinucleated myotube). The myotubes expressed myogenin, desmin, actin, and α-actinin (**Fig. 2B**), which are relevant markers for myotube differentiation. To estimate the differentiation potential, we next evaluated the efficiency of cell fusion, by staining of muscle cytoskeletal marker desmin and DAPI. We calculated a fusion index of 70.64% (SD 2.37), 69.32% (SD 3.73), 73.60% (SD 5.84), 68.94% (SD 9.17), 71.20% (SD 11.74) and 79.68% (SD 2.99) for donor 1, 2, 3, 4, 5 and 6, respectively, indicating a robust differentiation potential (**Fig. 2C**). Lastly, after prolonged culture (14 days) the cells formed sarcomeric structures, visible as striation in stainings for muscle cytoskeletal proteins α-actinin and desmin (**Fig. 2D**).

**Fig. 2:**
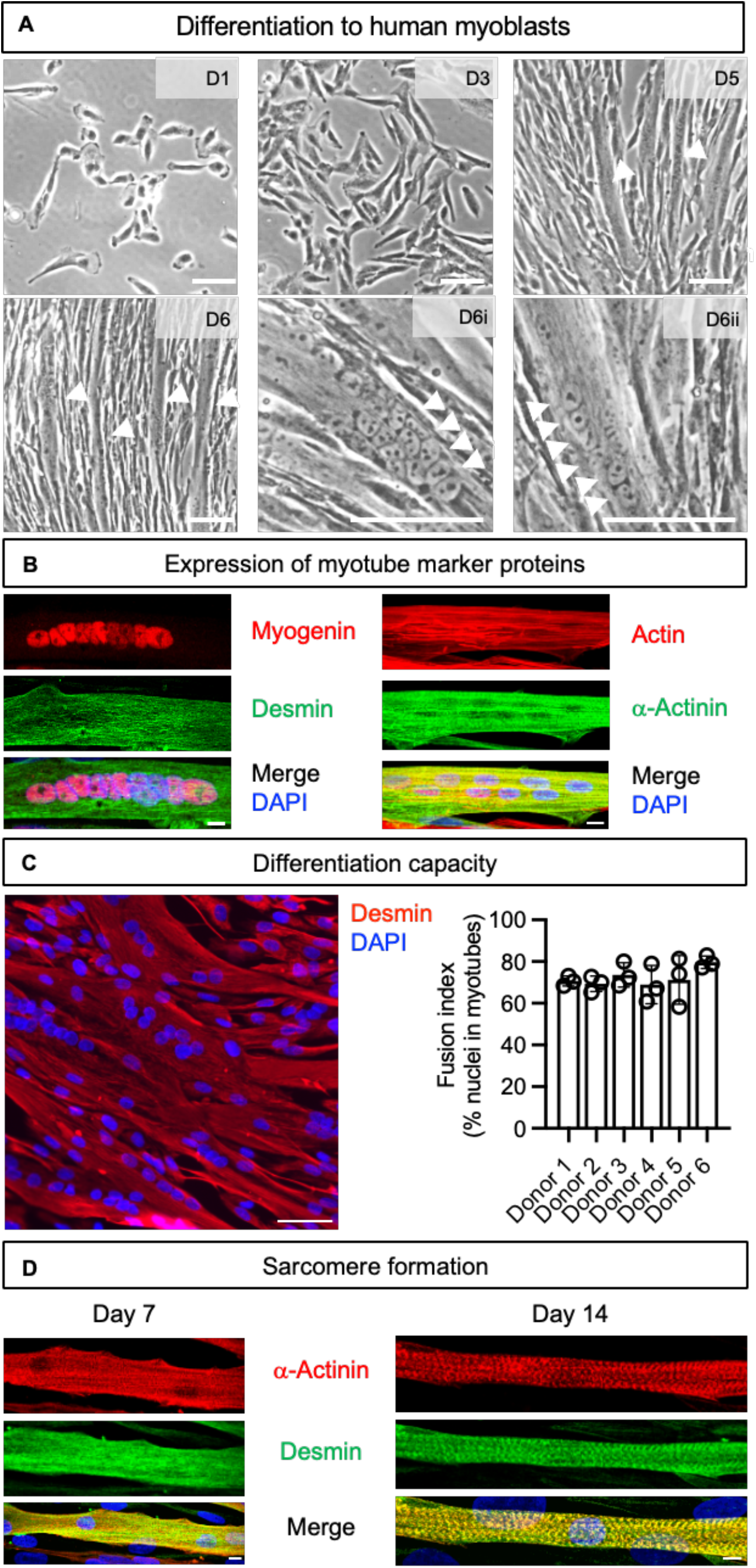
Isolated muscle stem cells differentiate to multinucleated myotubes. **(A)** Representative bright field microscopy images showing isolated myoblasts from donor 6 on day 1, 3, 5 and 6 (D1-D6) after seeding. The presence of myotubes observed on day 5 and 6 is indicated by arrowheads. Multiple nuclei per myotubes are indicated by arrowheads in (D6i) and (D6ii). The scale bars indicate 100 µm. **(B)** The myotubes expressed myotube marker proteins myogenin, actin, α-actinin and desmin. The scale bar indicates 10 µm. **(C)** The differentiation capacity of the muscle cells was determined in desmin/DAPI counterstaining, example image from donor 6. The scale bar indicates 50µm. The fusion index was determined as % of nuclei in multinucleated myotubes (≥2 nuclei/myotube, evaluation of ten microscopic images, N=3). **(D)** After 14 days in culture, the myotubes matured and developed sarcomeres, visible as striation pattern in desmin/α-actinin staining. The scale bar indicates 10 µm.

Stimulation of aneurally cultured myotubes with neural agrin can induce postsynaptic development and AChR clustering, thus providing an *in vitro* model of the postsynaptic part of the NMJ (*4, 50, 51*), contributing to our understanding of synaptic development (*10, 52*) and pathogenic mechanisms of MG autoantibodies at the NMJ (*1, 21, 22, 53–55*). We examined protein expression of key components of the agrin-LRP4-MuSK-DOK7 signal transduction pathway (schematic representation in **Fig. 3A**) and observed, as expected, expression and upregulation of DOK7, rapsyn, and AChRα upon differentiation to myotubes and following agrin stimulation (**Fig. 3B**, **Fig. S1**). Interestingly, LRP4 showed a downregulation upon differentiation by 63.09% when unstimulated and 53.30% when stimulated with agrin, while MuSK levels remained similar throughout differentiation. Importantly, AChR clustering could be induced on myotubes by stimulation with neural agrin and visualized by red-fluorescent α-bungarotoxin (**Fig. 3C-D**). While an increased AChR clustering upon agrin stimulation was observed for multiple passages, a decline of AChR cluster formation compared to the spontaneous AChR cluster formation of unstimulated cells occurred after passage 9 (**Fig. S2**). Therefore, for experiments requiring AChR cluster formation, we only used cells prior to passage 9. AChR clusters are henceforth referred to as *in vitro* NMJs. In the next step, we wanted to evaluate the model’s usefulness to study MG autoantibody binding and pathogenicity.

**Fig. 3:**
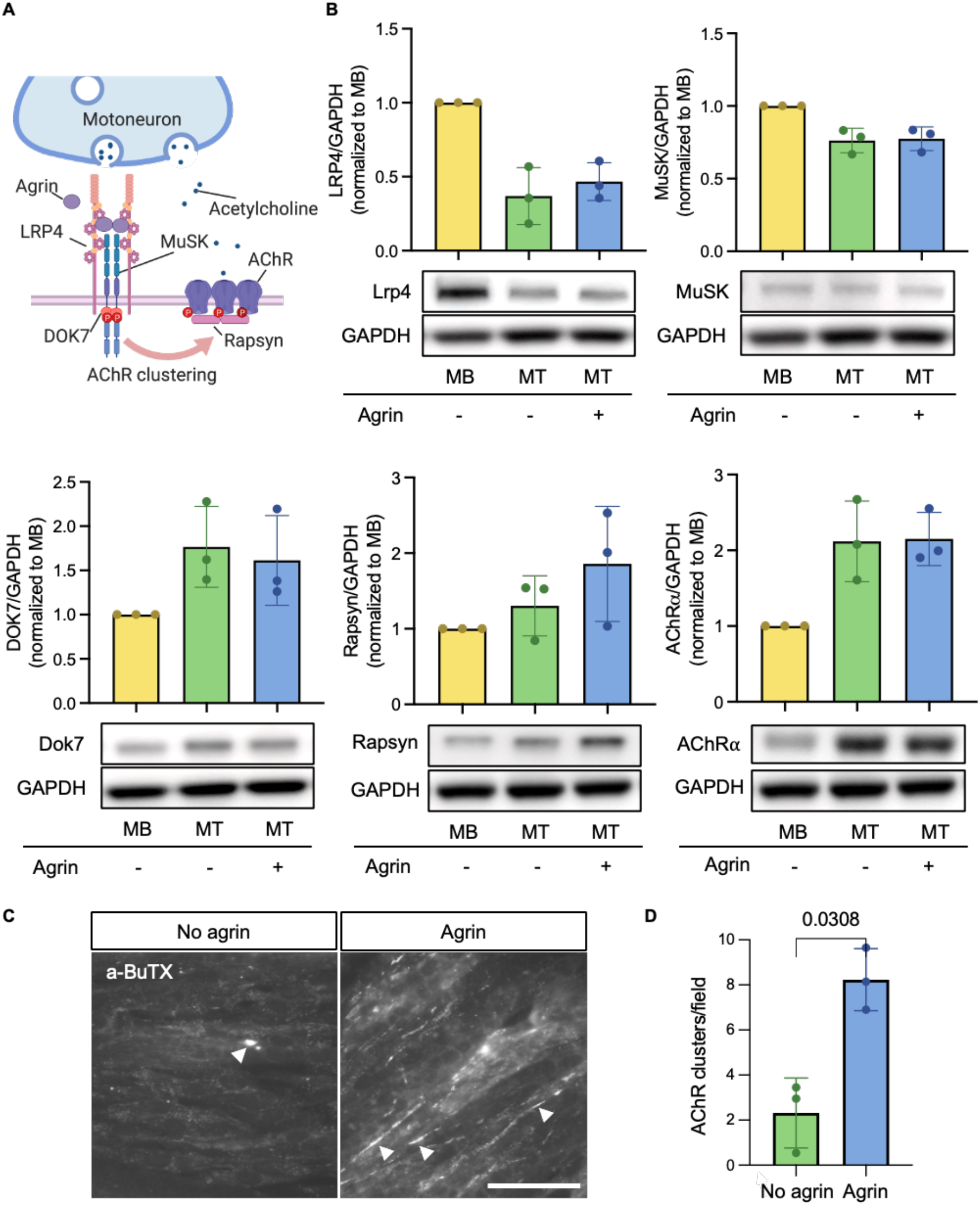
Human myotubes express neuromuscular proteins, downregulate LRP4 upon differentiation and are capable to cluster AChRs upon agrin stimulation. **(A)** Schematic representation of the postsynaptic neuromuscular junction. **(B)** Expression of LRP4, MuSK, DOK7, rapsyn and AChRα in myoblasts (MB) and myotubes (MT, without or with agrin stimulation) was evaluated by Western blot; normalized to MB condition; bands normalized to GAPDH **(C)** Representative immunofluorescence images of primary human muscle cells (passage 7) with and without agrin stimulation 16 h prior to α-bungarotoxin-AF594 AChR cluster staining. The formed AChR clusters are indicated by white arrowheads. The scale bar indicates 50 µm. **(D)** Quantification of 20 microscopic images per condition (mean and SD). Paired t-test. N=3.

### AChR- and MuSK-MG antibodies bind to and reduce AChR clusters in agrin-stimulated myotubes

To evaluate binding of patient autoantibodies to the *in vitro* NMJs, human myoblasts were differentiated to myotubes, and AChR clustering was induced by stimulation with neural agrin before incubation with healthy control or MG patient serum. AChRs were visualized with red-fluorescent α-bungarotoxin, and autoantibody binding with a green-fluorescent anti-human IgG secondary antibody. Colocalization of signal for human IgG and *in vitro* postsynaptic NMJs was considered an indication for autoantibody binding to AChR or MuSK, respectively (**Fig. 4**, **Table S2**). A pronounced binding of antibodies to the *in vitro* NMJs was observed in the majority of MG sera. The assays showed a sensitivity of 96.43% for MG antibodies (95%CI: 0.817-0.999, 27/28 MG sera positive, p<0.0001, Fisher’s exact test, Clopper–Pearson method), with 21/22 AChR-MG sera (sensitivity: 95.45%, 95% CI:0.772-0.999), and 6/6 MuSK-MG sera (100%, 95% CI:0.541-1.000) showing colocalization of patient IgG with in vitro NMJs.

**Fig. 4:**
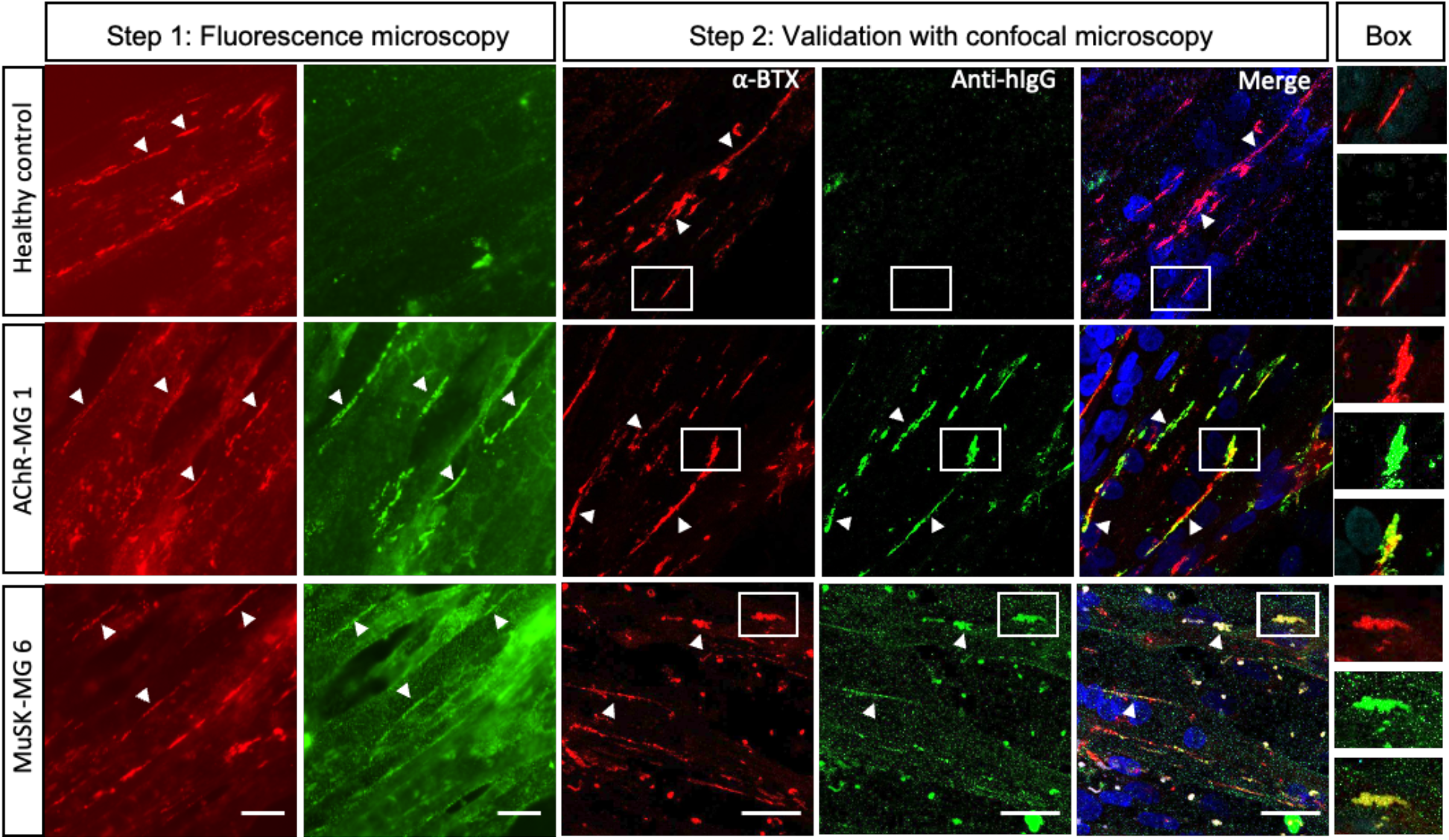
Muscle cell-based assay to detect NMJ autoantibodies. Colocalization of human IgG (anti-human IgG-AF488, Anti-hIgG, green fluorescence) with *in vitro* NMJs (AChR clusters, α-bungarotoxin-AF594, α-BTX, red fluorescence) is first identified by fluorescence microscopy (step 1), then validated in a second step by confocal microscopy. Arrowheads indicate *in vitro* NMJs (AChR clusters) and/or colocalized anti-hIgG binding. Representative images from a healthy control, AChR-MG and MuSK-MG patient. Box: indicating box selection from confocal images. Blue fluorescence: DAPI. Scale bars indicate 20µm.

Whenever the colocalization appeared less pronounced, which was the case with MuSK-MG sera, it was validated in a second step using confocal microscopy, (Fig. 4, bottom row). We thus provide proof-of principle data indicating that the human muscle cell-based assay is suitable to detect neuromuscular autoantibodies with high sensitivity and specificity.

As MG autoantibodies reduce AChR densities, e.g. by cross-linking and endocytosis of AChRs or by interference with MuSK activation (*22, 53, 56*), we tested the suitability of our model to study the effect of AChR- and MuSK autoantibodies on agrin-induced AChR clustering. We used experimental conditions as described previously in C2C12 mouse myotubes (*22, 57*) and incubated human myotubes for 16 hours with agrin and with or without serum from healthy controls or patients with AChR-MG or MuSK-MG. The AChR clusters were visualized with red-fluorescent α-bungarotoxin and quantified in a blinded manner from 20 microscopic fields (**Fig. 5**). AChR counts were normalized to the average number of AChR clusters of agrin-only-treated cells. Sera from MG patients reduced the average number of AChR clusters by 50.58-84.71% (AChR-MG 1: 34.65% of agrin (95%CI:6.51-62.79, p<0.0001); AChR-MG 3: 15.29% of agrin (95%CI: 0.55-30.03, p<0.0001); AChR-MG 22: 49.42% of agrin, (95%CI:6.47-92.37, p=0.0017); MuSK-MG 1: 37.95% of agrin (95%CI:26.89-49.01, p<0.0001); MuSK-MG 5: 34.32% of agrin (95%CI:19.24-49.39, p<0.0001); MuSK-MG 6: 31.49% of agrin (95%CI:8.65-54.32, p<0.0001) compared to the agrin-only control. Healthy control serum (pooled HC: 91.11%, 95%CI:76.51-105.71) reduced the AChR formation by only 8.89%; One-way ANOVA; multiple comparisons corrected with Dunnett’s post hoc test. Thus, the human muscle cell model presented herein is suitable for studying functional effects of MG antibodies on the AChR cluster formation as a pathogenic mechanism.

**Fig. 5:**
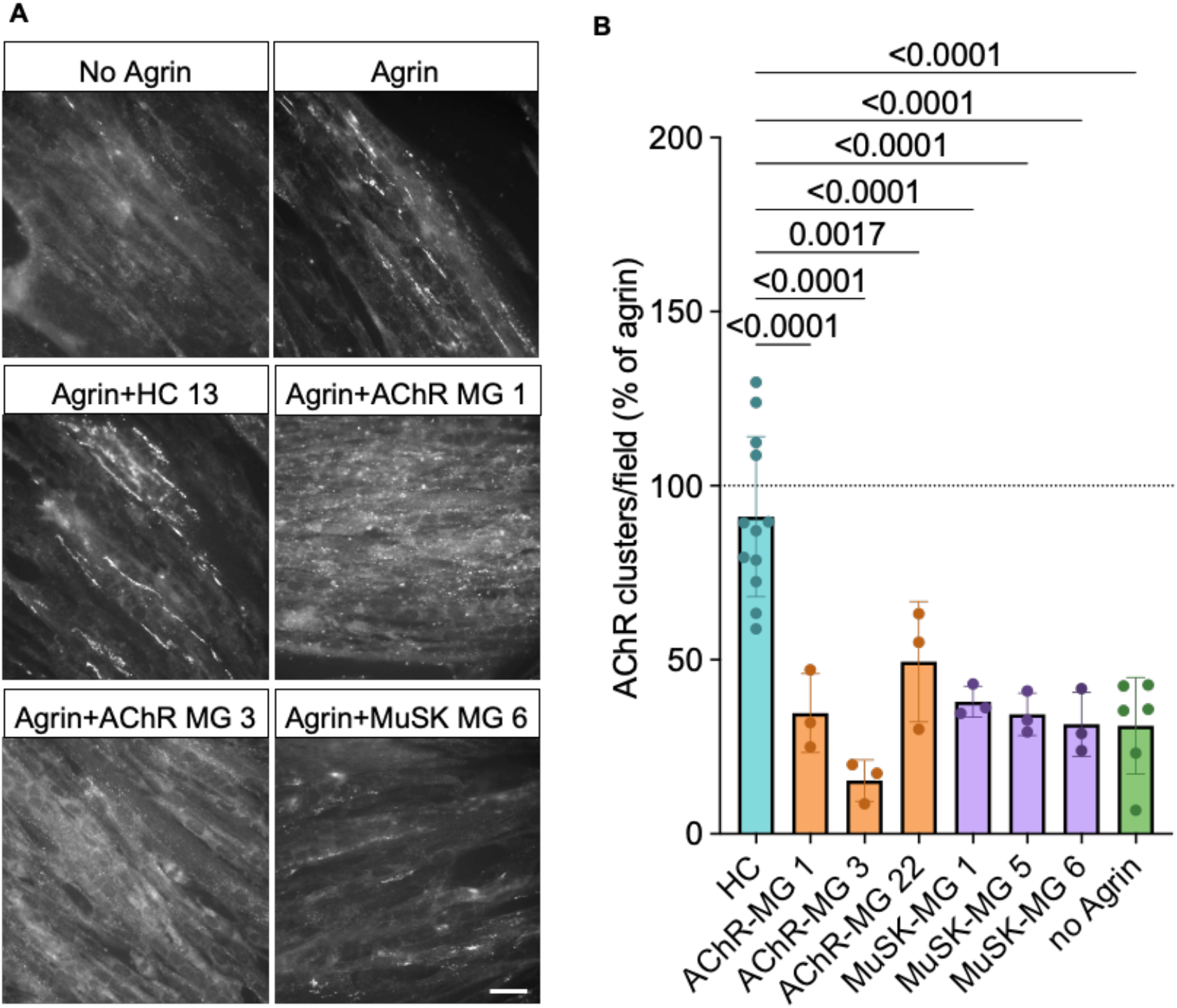
MG antibodies impair AChR clustering in primary human muscle cells. **(A)** Differentiated human myotubes were incubated 16h with agrin or no agrin, or agrin and serum from 4 healthy controls, 3 AChR-MG or 3 MuSK MG patients. AChR clusters were visualized by α-bungarotoxin-AF594. Scale bar indicates 20µm. **(B)** 20 microscopic fields with myotubes were selected in bright-field and fluorescent images acquired at 20x magnifications. AChR clusters ≥5µm were quantified under blinded conditions. The numbers of AChR clusters per microscopic field were normalized to AChR clusters in agrin only treated cells. N=3 for AChR-MG and MuSK-MG, N=4 for HC (pooled), N=6 for agrin-only and no agrin control and differentiation medium control. Ordinary one-way ANOVA, Dunnett’s post hoc test for multiple comparisons

## DISCUSSION

Despite major advances in the identification of pathogenic autoantibodies in MG, current diagnostic and experimental approaches are limited by antigen-specific assays and by the lack of accessible, physiologically relevant human models to study NMJ autoimmunity. To address this gap, we developed a human muscle cell model suitable for the study of MG antibodies, which combines physiological relevance with cost-efficiency and handling characteristics comparable to commonly used cell lines such as C2C12 or human rhabdomyosarcoma cells.

MG autoantibody diagnostics relies on the detection of antigen-specific autoantibodies, which is important for the stratification of MG patients, to guide treatment decisions and to further characterize patient autoantibodies and antigen-specific immune cells (*1*). There are however important limitations to this approach, including technical challenges of diagnostic assays, particularly for the detection of LRP4 autoantibodies (*28, 58*), and for patients with unknown antibodies. Intercostal muscle biopsies from SNMG patients showed IgG and complement deposition colocalized with the NMJ (*59*), suggesting the presence of pathogenic autoantibodies against yet undiscovered antigens. To further investigate this possibility, physiologically relevant *in vitro* models of NMJ autoimmunity are needed, which are accessible and easy to use. Human rhabdomyosarcoma cell lines such as TE671 (*41*) or CN21 (*60*) are frequently used in MG research (*40, 61, 62*), but are limited by expression of tumor-associated genetic alterations and lack of differentiation potential (*62*). Alternative models are based on C2C12 myoblasts, which are accessible, easy to handle, differentiate to myotubes and form AChR clusters (*22, 57, 63–65*), however, their murine origin (*34, 35*) confines translation to human disease because NMJ structure, gene expression and signaling pathways show species-specific differences (*36, 37*). Further, immortalization and prolonged culture of any cell lines may cause genomic abnormalities, changes in gene expression and function (*43, 66, 67*).

On the other hand, primary human muscle stem cells preserve the donor-specific genetic background, maintain the potential to self-renew, proliferate and differentiate into myogenic progenitor cells (*68–73*). They are suitable to study antibody effector functions such as complement activation (*54, 74*), but their widespread application has been limited by availability, cost and experimental flexibility (*75*).

Here, we propose a human muscle stem cell-based model that can be isolated with high purity from surgical or biopsy residual tissue, expanded, cryopreserved, and differentiated into myotubes, thereby enabling donor-specific human muscle models retaining key characteristics of the tissue of origin. Key components of agrin-induced AChR clustering pathway are expressed, including LRP4, MuSK, DOK7, rapsyn and AChRα, and we observed, as expected, an upregulation of DOK7, rapsyn and AChRα with differentiation (*6–9, 12, 15, 76*). Interestingly, LRP4 expression decreased during human myogenic differentiation, contrasting previous observations in murine C2C12 cells (*6*) and underlining the need to further investigate species-specific functions of LRP4 in human muscle biology.

Accordingly, the herein presented cell-based assay provides the basis for a complementary diagnostic approach that overcomes current limitations of antigen-specific assays in MG. Both AChR- and MuSK-antibody-positive sera demonstrated IgG binding to *in vitro* NMJs. In future studies, this assay can be used as basis for the detection and characterization of postsynaptic NMJ antibodies and to support efforts in antigen discovery. Further, as sera from patients with AChR or MuSK antibodies reduced AChR clusters in culture, we suggest our model is useful as a basis for functional assays, to characterize the effect of MG antibodies on the human agrin-LRP4-MuSK-DOK7 signal transduction pathway (*1*) and to study complement activation at the NMJ *in vitro* (*54*).

Beyond autoimmune MG, we previously used cells from donor 4 and donor 6 to study the effects of plectin ablation on cytoskeletal architecture (*77*). Plecstatin-1-treatment of differentiated myotubes resulted in increased bundling of desmin and vimentin, which was reminiscent of desmin-rich protein aggregates in muscle biopsies from patients with epidermolysis bullosa simplex with muscular dystrophy (*1, 2, 78*). The model may thus also enable studies of genetic NMJ disorders, including congenital myasthenic syndromes, by providing a human muscle background suitable for investigating disease-associated molecular defects.

### Limitations

As expected in primary cells, after prolonged passaging we observe a phenotypic drift, which manifested as loss of AChR clustering potential, suggesting the need to use early passages of cells. However, the proposed culturing strategy allows for a high yield of cells at early passages, which can be cryopreserved and allow long-term supply with relevant cells. This approach is similar to a previously reported strategy to obtain high numbers of low-passage C2C12 cells (*79*). Further, the aneural culture model lacks complexity as the presynaptic parts of the NMJ, including motor neurons and Schwann cells are missing, but the cells provide a basis to develop more complex co-culture systems in the future. Taken together, depending on the scientific question, our aneural culture model is a highly feasible and important new tool in MG research.

### Conclusions

We present a simple and cost-efficient method to isolate primary human muscle cells with a high yield of early passage cells that can be cryopreserved, cultured and differentiated into myotubes. These muscle cells are the basis for a new human *in vitro* model of the human NMJ, a novel muscle CBA for detection of neuromuscular autoantibodies and for functional assays to study effects of MG antibodies on agrin-induced AChR clustering *in vitro*. Our model will thus provide the basis for functional studies of autoimmune and genetic diseases of the muscle and the NMJ.

## MATERIALS AND METHODS

### Muscle tissue and ethical considerations

Adult human muscle was obtained from surgical waste from anterior cruciate ligament reconstruction surgery, removed from tendons of *musculus semitendinosus* and *musculus gracilis.* The study has ethical approval by the institutional review boards of the Medical University of Vienna, Austria (EK 1442/2017).

### Cohort

The cohort comprised of 22 AChR-MG patients (AChR-MG1 – AChR-MG22), 6 MuSK-MG patients (MuSK-MG1 – MuSK-MG6) and aged-matched 25 healthy controls (HC1 – HC25). 8 AChR-MG sera (AChR-MG 5-12) were recruited at the Charité Berlin. One sample was kindly provided by the Azienda Ospedaliero-Universitaria Pisana, Pisa, Italy. All other sera were stored in the biobank at the Division of Neuropathology and Neurochemistry of the Medical University of Vienna. Sera were collected from 2011-2025. Patient data such as age, sex, MGFA and age of onset are summarized in **table S2**.

### Isolation and maintenance of primary human muscle cells

Primary human muscle cells were isolated as described before (*46–49*). Briefly, human muscle tissue was collected in DPBS+P/S (DPBS, product number 14190136, Thermo Fisher Scientific, Waltham, MA, USA, supplemented with 1% penicillin/streptomycin, product number 15140122, Thermo Fisher Scientific, Waltham, MA, USA) at room temperature immediately after surgery. The isolation was started within six hours after collection. Muscle fragments were placed in a cell culture dish with DPBS+P/S and adipose and connective tissue were removed using a stereomicroscope (Stemi 508, Zeiss, Oberkochen, Germany). The remaining muscle tissue was then finely minced with the use of two scalpels and transferred to a 50mL tube containing 15mL of dissociation medium (DPBS supplemented with 25 mM glucose, 25 mM sucrose, 5.4 mM KCl, 0.05 mg/ml gentamicin (product number: G1397, Gibco, Thermo Fisher Scientific, Waltham, MA, USA), 2.5 mg/ml trypsin (Sigma-Aldrich, T4799), and 220 U/mL collagenase type II (product number: 17101-015, Gibco, Thermo Fisher Scientific, Waltham, MA, USA)) and incubated at 37°C with gentle agitation for 1 hour. The supernatant was collected and centrifuged for 5 min at 300 g at room temperature, and the pellet was resuspended in 10mL DPBS+P/S (first muscle cell suspension). The remaining muscle fragments were incubated for another hour with 15mL of fresh dissociation medium, followed by mechanical dissociation by trituration (10x using a 10mL serological pipette), then DPBS+P/S was added to 50mL followed by centrifugation for 5 min at 300 g. The pellet was resuspended in 10mL DPBS+P/S (second muscle cell suspension). Both muscle cell suspensions were combined and pipetted with pressure through a 100 µm cell strainer. To maximize the cell recovery, the strainer was rinsed 4–6 times with 5mL of DPBS+P/S. The cell suspension was centrifuged (300-400g, 5 min), and the resulting pellet was resuspended and cultured in muscle cell growth medium (Advanced DMEM/F12 (product number: 12634010, Gibco, Thermo Fisher Scientific, Waltham, MA, USA) supplemented with 20% FCS (product number: 10270106, Thermo Fisher Scientific, Waltham, MA, USA), 50 μg/ml fetuin (product number: F3385, Sigma Aldrich, St. Louis, MO, USA), 7 mM glucose, 4 mM L-glutamine (product number: 25030024, Gibco, Thermo Fisher Scientific, Waltham, MA, USA), 1% pen/strep (product number: 15140122, Gibco, Thermo Fisher Scientific, Waltham, MA, USA), 2.5 μg/ml amphotericin B (product number: 15290018, Gibco, Thermo Fisher Scientific, Waltham, MA, USA), 8 ng/ml FGF (product number: F0291, Sigma Aldrich, St. Louis, MO, USA), 20 ng/ml EGF (product number: SRP3027, Sigma Aldrich, St. Louis, MO, USA), 400 ng/ml dexamethasone (product number: D4902, Sigma Aldrich, St. Louis, MO, USA), and 200 ng/ml insulin (product number: I0516, Sigma Aldrich, St. Louis, MO, USA)) on Nunc™ cell culture dishes (product number: 150350, Thermo Fisher Scientific, Waltham, MA, USA) at 37°C and 5% CO_2_. The culture dish was monitored daily for cell attachment. In the initial days of culture, cellular debris from the isolation was visible, which was not always distinguishable from contamination, thus the cell culture dishes were kept in quarantine boxes. Six days post-isolation, the medium was changed to 10mL of fresh growth medium. Once adherent cells formed irregular patches (starting around day 10 post isolation), these were dissociated using trypsin-EDTA (product number: 25200056, Thermo Fisher Scientific, Waltham, MA, USA) and replated, with cell culture dish size adjusted to accommodate a cell density below 40%. When the cells reached 40% confluence in a 100 mm culture dish, the cells were defined as passage P0.

The muscle cell identity was validated by immunocytochemistry staining for desmin (described below) and the cells were henceforth referred to as myoblasts. For maintenance, myoblasts were cultured at densities of 800–3500 cells/cm² (for 1 to 3 days in culture) in muscle cell growth medium, and passaged at approximately 40% confluence. Cells were passaged latest every 3 days to avoid differentiation. The myoblasts were frozen in muscle cell growth medium supplemented with 10% DMSO (product number A3672,0100, AppliChem GmbH) and stored in liquid nitrogen.

### Human myotube differentiation and AChR clustering

For differentiation of myoblasts to myotubes, we seeded 3x10^4^ myoblasts per coverslip coated with 0.2% gelatine in 24-well plates in 500 µl of human muscle cell growth medium. The cells were incubated at 37°C 5% CO_2_ and left to proliferate, with medium exchange (growth medium) once 2 days after seeding if a confluence of 50% was not reached yet. When human myotubes were fully formed on day 6, the culture medium was replaced by differentiation medium (DMEM, high glucose, GlutaMAX™ Supplement (product number: 10566016, Thermo Fisher Scientific, Waltham, MA, USA), 5% horse serum (product number 16050-122, Gibco), 0.1µg/mL gentamicin (product number: G1397, Sigma Aldrich, St. Louis, MO, USA), 0.1µg/mL insulin (product number: I6634, Sigma Aldrich, St. Louis, MO, USA) supplemented with rat agrin (*51*), plasmid kindly gifted by Ruth Herbst and produced in HEK293T) and incubated for 16 h at 37°C 5% CO_2_ to allow for induction of AChR clusters. To analyze the pathogenic effects of MG patient serum on AChR clustering, agrin as well as healthy control or MG patient sera were added at 1:10 dilution to the differentiation medium. Next, the cells were incubated with α-bungarotoxin conjugated to AF594 (product number: B13423, Invitrogen, Thermo Fisher Scientific, Waltham, MA, USA) diluted 1:2000 in blocking medium I (DMEM supplemented with 1% BSA (product number: A8531 / A9647, Sigma Aldrich, St. Louis, MO, USA) and 20mM HEPES (CAS# 7365-45-9, Art# 9105.3, Carl Roth, Karlsruhe, Germany), sterile filtered) for one hour in the incubator at 37°C, 5% CO2 in the dark, washed gently 3x 10 minutes with DMEM (product number D6429, Sigma-Aldrich, St. Louis, MO, USA), at 37°C in the incubator, and once with PBS. Next, the coverslips were fixed in 4% paraformaldehyde, washed 3x with PBS, stained with DAPI, washed 3x with PBS and mounted with Aqua-Poly/Mount (product number: 18606-20, PolySciences, Warrington, PA, USA) before analysis by fluorescence microscopy (Olympus BX63, Olympus, Tokyo, Japan). To quantify AChR clustering, the samples were blinded throughout the experiment, and per condition 20 microscopic fields containing myotubes were selected in bright field at 20x magnifications, before microscopic images of red fluorescence and DAPI were acquired using cellSens image analysis software (Olympus, Tokyo, Japan) and analyzed for AChR cluster number and length using ImageJ software (Version 1.54k; National Institute of Mental Health, Bethesda, Maryland, USA). For prolonged differentiation (8-14 days), the cells were seeded directly on plastic to allow for a stronger attachment.

### Muscle cell-based assay

Human myoblasts were seeded, differentiated and stimulated with agrin as outlined above. To detect binding of patient autoantibodies, the myotubes were incubated with healthy control, AChR-MG or MuSK-MG sera diluted 1:50 - 1:100 and α-bungarotoxin conjugated to AF594 in blocking medium II (DMEM (product number: D6429, Thermo Fisher Scientific, Waltham, MA, USA), supplemented with 0.01 g/ml BSA (product number A9647, Sigma Aldrich, St. Louis, MO, USA), 20 mM HEPES (product number: 15630-056, Gibco, Thermo Fisher Scientific, Waltham, MA, USA) and 10% normal donkey serum (product number: S30, EMB Millipore, Darmstadt, Germany)) for 2 h at room temperature. The cells were washed 3x with medium for 10 min at 37°C and 5% CO_2._ Next, the cells were fixed with 4% PFA and washed 3 times with PBS. Human antibody binding to the muscle cells was visualized using Alexa Fluor 488 conjugated anti-human secondary antibody (product number: A-11013, Invitrogen, Thermo Fisher Scientific, Waltham, MA, USA) at a dilution of 1:750 at room temperature. After washing 3 times with PBS the nuclei were stained with DAPI and mounted, and colocalization of human antibodies to the AChR clusters was evaluated by fluorescence microscopy (Olympus BX63, Olympus, Tokyo, Japan). Where necessary, the colocalization was validated by confocal microscopy at 40x and 60x magnification (Leica Stellaris 8 laser scanning microscope, Leica, Wetzlar, Germany).

### Immunocytochemistry

Each 2 x 10^4^ or 3 x 10^4^ (for differentiation) primary human muscle cells or 1-3x10^5^ HEK293 cells were seeded per glass coverslip coated with 0.01% Poly-L-Lysine (product number: P2636, Sigma Aldrich, St. Louis, MO, USA, for HEK293) or 0.2% gelatine (primary muscle cells) per well in a 24 well-plate. 3 days after seeding, the myoblasts were fixed for 5 minutes with 4% paraformaldehyde at RT. For cytoskeletal stainings, myoblasts were differentiated to myotubes as outlined above. The cells were 3x washed with PBS, then incubated with PBS supplemented with 0.3% TritonTM X-100 for 5 min at RT, washed 3x with wash medium (DMEM, product number: D6429, Thermo Fisher Scientific, Waltham, MA, USA) and incubated with 1:100 anti-desmin antibody (product number: M0760, Agilent Technologies Dako, Glostrup, Denmark, diluted in blocking medium II (DMEM (product number: D6429, Thermo Fisher Scientific, Waltham, MA, USA), supplemented with 0.01 g/ml BSA (product number A9647, Sigma Aldrich, St. Louis, MO, USA), 20 mM HEPES (product number: 15630-056, Gibco, Thermo Fisher Scientific, Waltham, MA, USA) and 10% normal donkey serum (product number: S30, EMB Millipore, Darmstadt, Germany)) for 30-minutes at 37°C. Antibodies used for characterization of myotubes acquired with confocal microscopy were 1:100 Desmin CT-1 (*80*), anti-Myogenin (Dako, clone F5D, Santa Clara, USA), 1:1000 anti-actin (product number: PF00003, Proteintech Group, Inc., Rosemont, IL, USA), 1:700 anti-actinin (product number: A7811, Sigma Aldrich, St. Louis, MO, USA). The cells were washed 3x for 10 minutes with medium at 37°C, 5% CO_2_, and incubated with Alexa Fluor 488- or 594-conjugated goat anti-mouse secondary antibody (product number: A11001, Invitrogen, Thermo Fisher Scientific, Waltham, MA, USA) diluted 1:750 in blocking medium I. The cells were washed, stained with 2 µg/ml DAPI (product number: 10236276001, Sigma-Aldrich, St. Louis, MO, USA) and mounted with Aqua-Poly/Mount mounting medium (product number: 18606-20, PolySciences, Warrington, PA, USA) before microscopy analysis with an Olympus BX63 and CellSens image analysis software (Olympus, Tokyo, Japan), and nuclei in single cells and multinucleated myotubes were quantified from 10 microscopic images per condition using ImageJ software. The fusion index was calculated as the percentage of nuclei in myotubes, obtained from 10 images at 20x magnification per condition.

### Western blot

Primary human myoblasts and myotubes were washed with cold PBS, and lysed with RIPA lysis buffer (product number: 20-188, Millipore Sigma, Burlington, MA, USA) supplemented with protease inhibitor cocktail (product number: P8340, Sigma Aldrich, St. Louis, MO, USA). The cells were gently agitated for 15-60 min at 4 °C and subsequently harvested using a cell scraper (product number: 3010, Corning® Small Cell Scraper, Corning, NY, USA). The lysates were transferred to 1.5 mL tubes and centrifuged at 15,294 g for 15 min at 4 °C. The supernatants were collected and stored at -20 °C until further use. The protein concentration was measured using the Pierce™ BCA Protein Assay Kits (product number: 23225, Thermo Fisher Scientific, Waltham, MA, USA) and proteins were separated on NuPAGE™ Bis-Tris 4–12% Mini Protein Gels (product number: NP0335BOX, Thermo Fisher Scientific, Waltham, MA, USA) with Precision Plus Protein Dual Color standards (product number: 1610374, Bio-Rad Laboratories, Hercules, CA, USA) followed by transfer on PVDF membranes using a semi-dry transfer system (Invitrogen^TM^ Power Blotter, Thermo Fisher Scientific, Waltham, MA, USA) at the mixed molecular weight (MW) setting for proteins of 25-150 kDa (1.3 A, 7 min) or the High MW setting for proteins above 150 kDa (1.3 A, 10 min). Proteins were detected using 1:200 mouse anti-LRP4 (product number: B258235, BioLegend, San Diego, CA, USA), goat anti-DOK7 (product number AF6398, R&D Systems, Minneapolis, MN, USA), 1:2000 rabbit anti-rapsyn (product number: EPR9759, Abcam, Cambridge, UK), 1:500 goat anti-human MuSK (product number AF3904, R&D Systems, Minneapolis, USA), 1:500 mouse anti-MyoD (product number: MA1-41017, Invitrogen, Thermo Fisher Scientific, Waltham, MA, USA), mouse anti-AChRα (product number: 610988, BD Biosciences, San Jose, CA, USA) or 1:10,000 mouse anti-GAPDH (product number: SC-32233, Santa Cruz Biotechnology, Dallas, USA) diluted in EveryBlot Blocking Buffer (product number: 12010020, Biorad). As secondary antibodies, 1:10,000 donkey anti goat IgG HRP (product number: AB 234O390, Lot number: 177481, Jackson ImmunoResearch Laboratories, West Grove, USA), 1:1000 anti-mouse IgG HRP (product number: P0447, Agilent Dako, Santa Clara, USA) or 1:2000 goat anti-rabbit IgG HRP (product number: P0448, Agilent Dako, Santa Clara, USA) were used with BioRad Clarity™ Western ECL Substrate (product number: 170-5061, Biorad, Hercules, CA, USA) and detected with a Fusion FX imaging system (Vilber Bio Imaging SAS, Collégien, France). Quantification of intensity was performed with the Evolution-Capt Edge software. For background subtraction, the rolling ball function was used. Intensities were normalized to the intensity of the myoblast band.

### Statistical analysis

Data are presented as mean ± standard deviation (SD). Normality was assessed using the Shapiro-Wilk test. If the test indicated a deviation from normal distribution (p < 0.05), data were considered non-normally distributed. Depending on the distribution, statistical analysis was performed using one-way ANOVA with Dunnett’s post hoc test for normally distributed data or the Kruskal–Wallis test with Dunn’s post hoc test for non-normally distributed data. Statistical significance was defined as a two-tailed p-value ≤ 0.05. Sensitivity and specificity were calculated with 95% confidence intervals using Fisher’s exact test and Clopper–Pearson method. All analyses were performed using GraphPad Prism version 9.1.0 (GraphPad Software, San Diego, CA, USA).

## List of Supplementary Materials

Table S1: Demographic data on muscle cell donors

Table S2: Demographic and clinical data on AChR-MG, MuSK-MG and healthy control sera donors

Fig. S1: Longitudinal analysis of NMJ protein expression.

Fig. S2: Longitudinal analysis of agrin-induced AChR cluster formation.

Fig. S3: AChR- and MuSK-MG sera show antibody binding in proximity to AChR clusters.

Fig. S4: Validation of antibody binding to in vitro NMJs.

## Acknowledgments

We thank Birgit Niederreiter, Benni Koch, Andreas Bugelnig, Verena Endmayr and Carmen Haider for technical assistance. We thank Roberta Ricciardi for her collaboration.

## Author contributions

Conceptualization: IK, LWi, MW, SHo,

Methodology: IK, LWe, LWi, MW

Investigation: MW, ASZ, KS, TD, OK, AW, RW, MR, PD, NM, KT, MP, PF, CE, FF, MM, HC, RHo, SH, RHe, CA, RHe, FZ, LWe, LWi, IK

Visualization: MW, IK, PD, LWi

Funding acquisition: IK, Sho

Project administration: IK

Supervision: IK

Writing – original draft: IK, MW

Writing – review & editing: MW, ASZ, KS, TD, OK, AW, RW, MR, PD, NM, KT, MP, PF, CE, FF, MM, HC, RHo, SH, RHe, CA, RHe, FZ, LWe, LWi, IK

## Data and materials availability

All data generated or analyzed during this study are available from the authors upon reasonable request. Information regarding the source of the materials used in this study is provided in the main manuscript.

