## SUPPLEMENTARY MATERIALS for "A primary human muscle cell-based assay for detecting myasthenia gravis autoantibody binding and assessing AChR cluster impairment"

### Characteristics of study participants (muscle tissue donors)

Human muscle was obtained from surgical waste from anterior cruciate ligament reconstruction surgery of six otherwise healthy human donors. The demographic information of the donors is available in **Table S1**.

**Supplementary table S1: Demographic data on muscle cell donors**

| Donor number | Age at sample | Sex (F = female, M = male) |
|---|---|---|
| D1 | 56 | M |
| D2 | 54 | F |
| D3 | 34 | F |
| D4 | 31 | M |
| D5 | 25 | M |
| D6 | 20 | M |
| Median | 32·5 years | 33·3% F |

### Characteristics of study participants (serum donors)

Demographic characteristics of study participants and serum binding properties of 25 healthy individuals, 22 AChR-MG and 6 MuSK-MG patients used in the study (**Table S2**).

**Supplementary table S2: Demographic and clinical data on AChR-MG, MuSK-MG and healthy control sera donors**

| Characteristics | Healthy donors (n=25) | AChR-MG (n=22) | MuSK-MG (n=6) |
|---|---|---|---|
| Age, median (IQR), in years | 45 (33·5-58) | 47 (34·25-75) | 37 (22·5-52·5) |
| Female, n (%) | 19 (76) | 7 (31·82) | 4 (66·7) |
| Male, n (%) | 6 (24) | 15 (68·18) | 2 (33·3) |
| MGFA scores | | | |
| MGFA score 0, n (%) | - | 1 (4·5) | 1 (16·7) |
| MGFA score I, n (%) | - | 5 (22·7) | 1 (16·7) |
| MGFA score II, n (%) | - | 11 (50) | 2 (33·3) |
| MGFA score III, n (%) | - | 0 (0) | 2 (33·3) |
| MGFA score IV, n (%) | - | 0 (0) | 0 (0) |
| MGFA score unknown, n (%) | - | 5 (22·7) | 0(0) |
| Age of onset <50, n (%) | - | 12 (54·5) | 5 (83·3) |
| Age of onset >50, n (%) | - | 7 (31·8) | 1 (16·7) |
| Age of onset unknown, n (%) | - | 3 (13·6) | - |
| Binding in proximity of AChR clusters | 0/25 (0%) | 21/22 (95·5%) | 6/6 (100%) |

### Longitudinal expression of NMJ proteins in primary human muscle cells

Expression of key NMJ proteins in myoblasts (MB) and myotubes (MT) over time was evaluated by Western blot (**Fig. S1**). We observed that protein expression levels were similar between P5-P12, with a trend for increased Lrp4 and Rapsyn expression in passage P12.

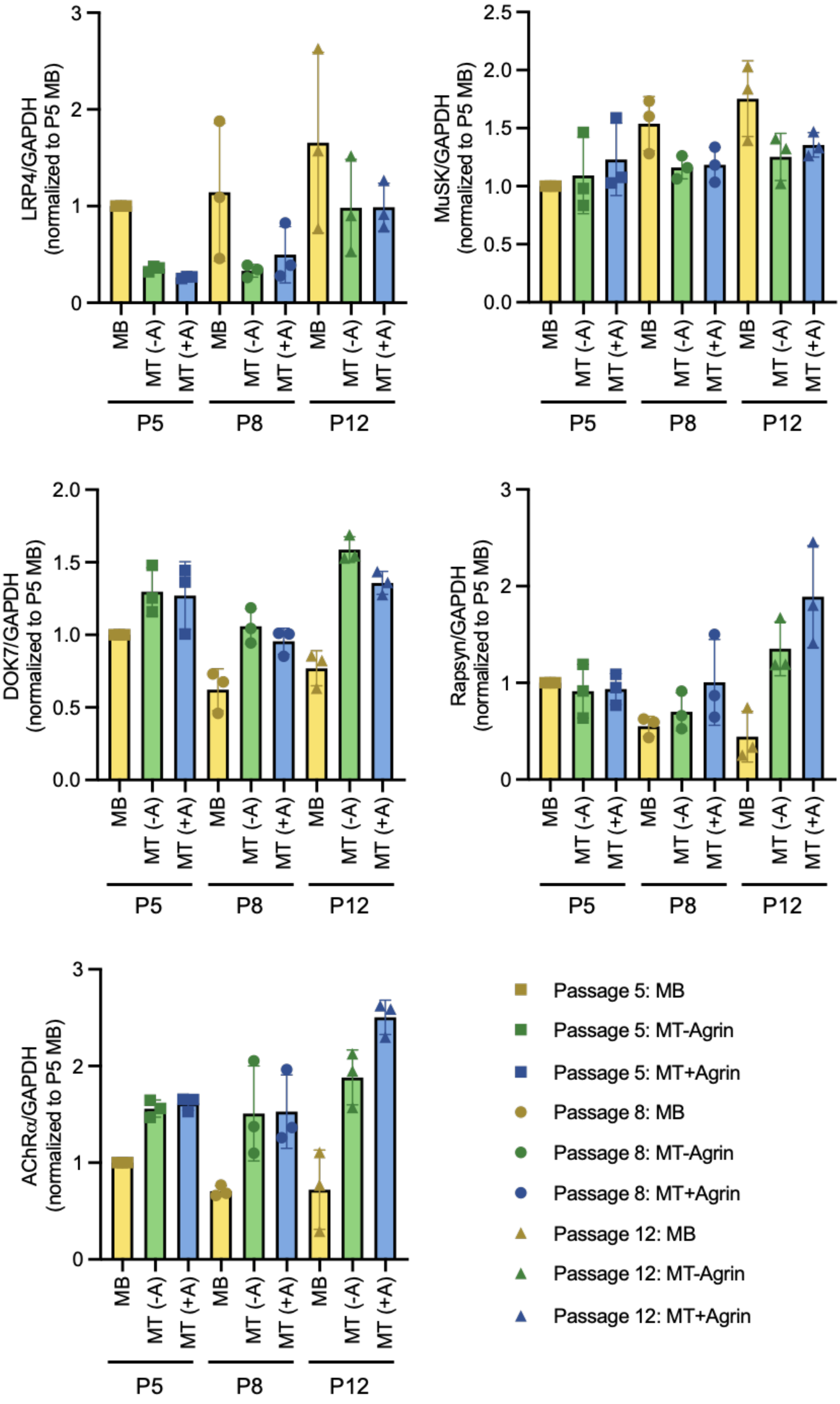


**Supplementary fig. S1: Longitudinal analysis of NMJ protein expression.** Expression levels of Lrp4, MuSK, Dok7, Rapsyn and AChR for cell lysates of myoblasts and myotubes from passage 5, 8 and 12 was determined by Western blot. Bands are normalized to the expression of GAPDH. Data is normalized to P5 myoblasts; N=3 (technical replicates)

### Longitudinal analysis of AChR cluster formation upon agrin stimulation compared unstimulated myotubes

We evaluated the cell's ability to maintain AChR clustering potential over time (**Fig. S2**). We observed that long term culturing affected AChR clustering potential, suggesting use of cells prior to P10.

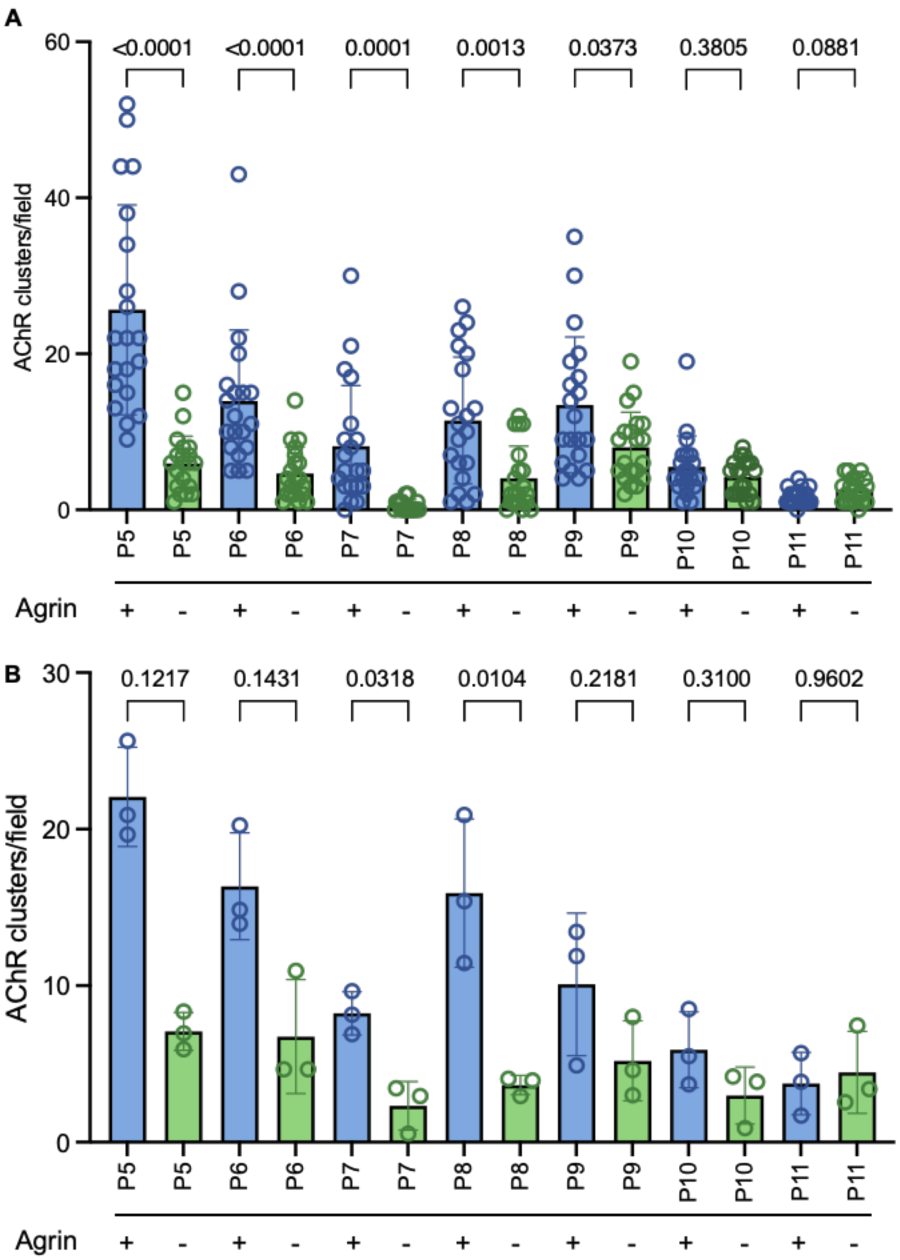


**Supplementary fig. S2: Longitudinal analysis of agrin-induced AChR cluster formation.** We evaluated AChR clusters ≥ 5 µm in cells with and without agrin stimulation between P5 and P11. Stimulation with agrin significantly increased the number of AChR clustering up until P9. **(A)** Representative data from n=1 for P5-P11. High variability between cluster counts from individual pictures can be observed. Agrin significantly induced AChR cluster formation until P9. Differences between the stimulated and unstimulated condition was analyzed by Mann-Whitney test. **(B)** Pooled data from three experiments did not reach significance for all passages due to inter-experimental variability. Differences between stimulated and unstimulated conditions were analyzed by Kruskal-Wallis test; N=3

### Colocalization of antibodies to the in-vitro postsynaptic NMJ was detected in 96·43% of positive controls in the muscle cell-based assay

Data for all patients from serum staining in the muscle CBA, not previously shown in **Fig. 2** of the main manuscript, are presented in **Fig. S3**. 21/22 (95·45%) of AChR-MG showed colocalization of antibodies to AChR clusters in this assay. AChR-MG 10 showed no colocalization to the AChR clusters. 6/6 (100%) of tested MuSK-MG sera contained antibodies binding in proximity to the *in vitro* NMJ. As MuSK antibodies are not expected to bind directly to the AChR, but in proximity, a much less clear staining is observable. Therefore, we analyzed the MuSK-MG cases with confocal microscopy (**Fig. S4**)

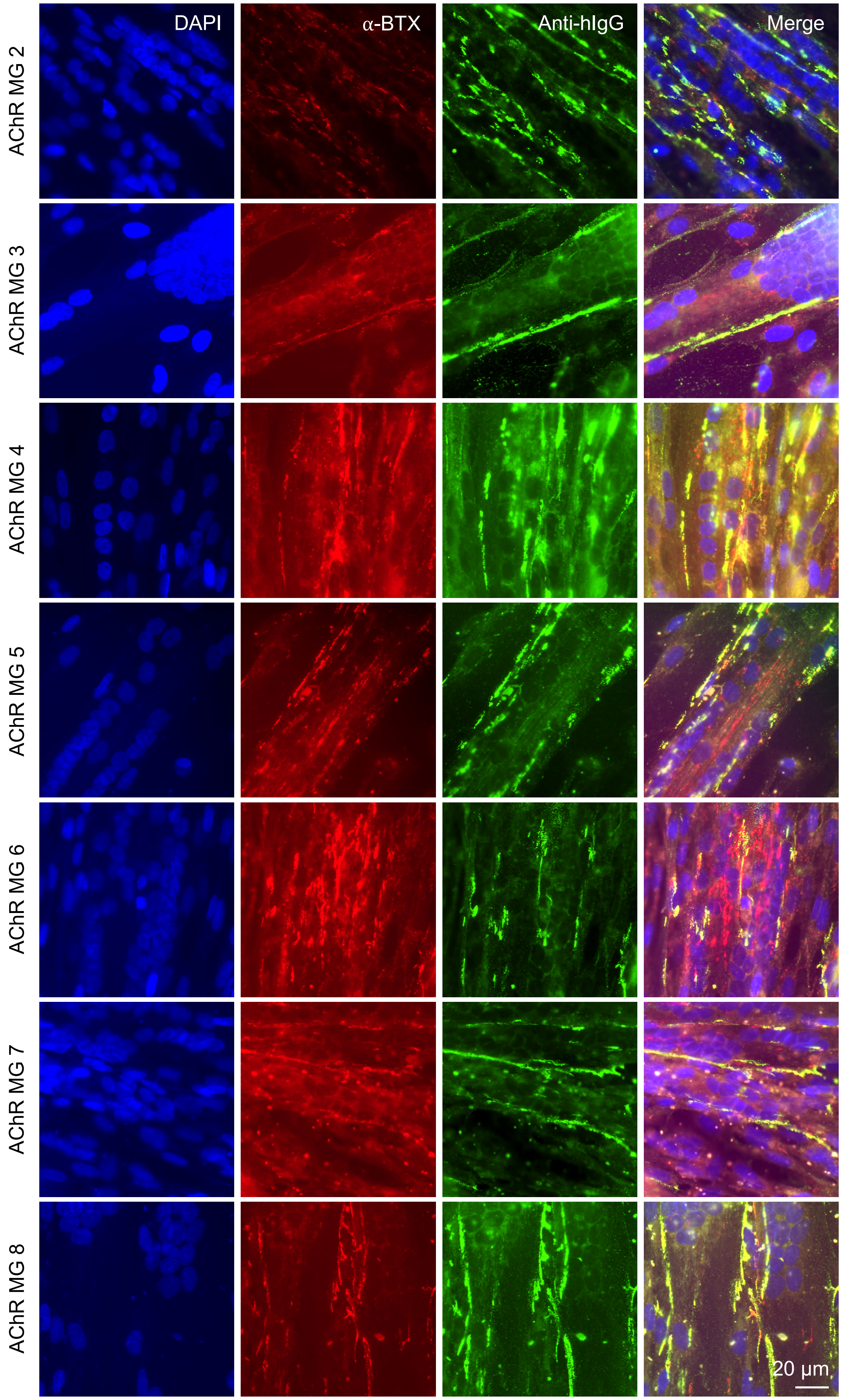




Continuation next page

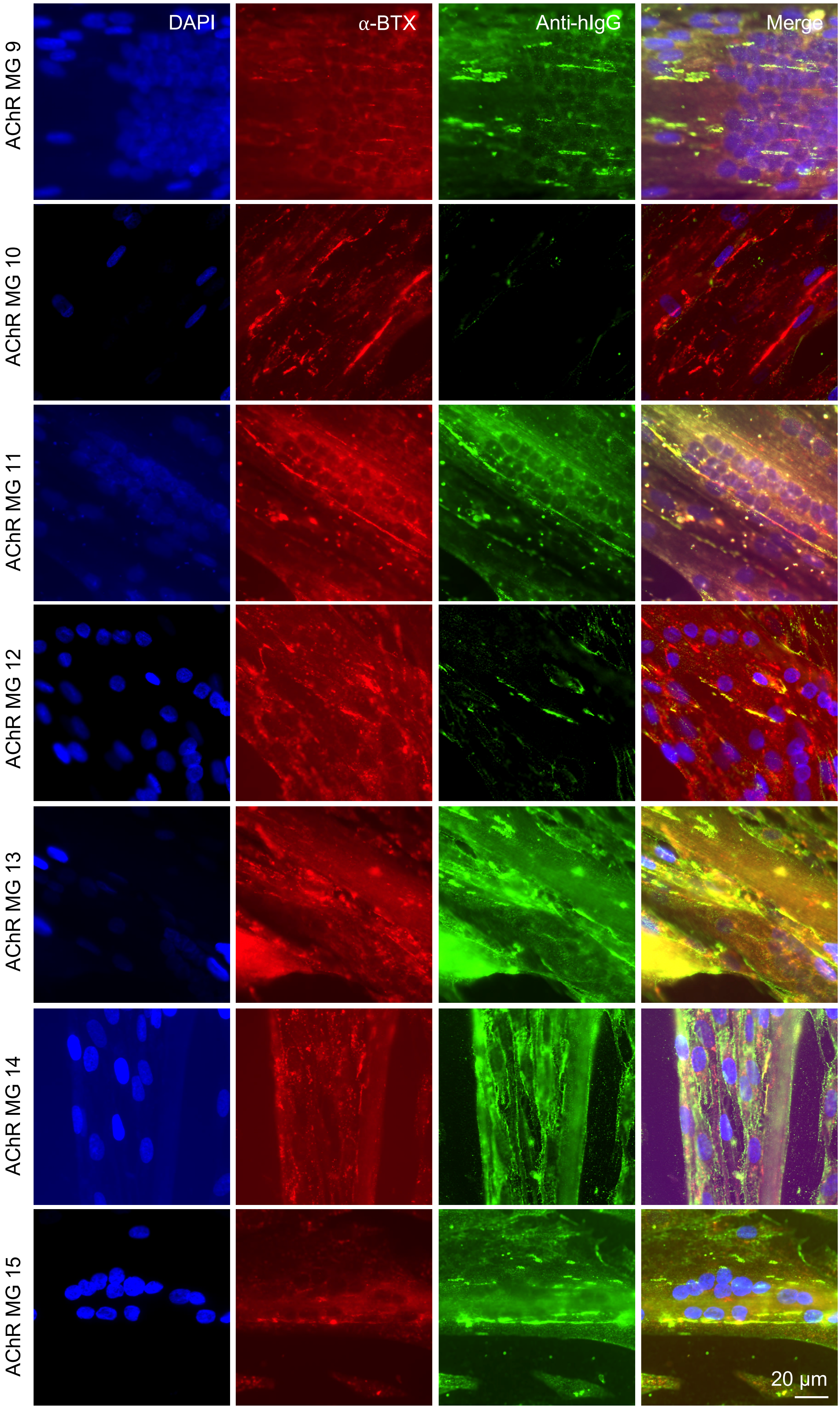


Continuation next page

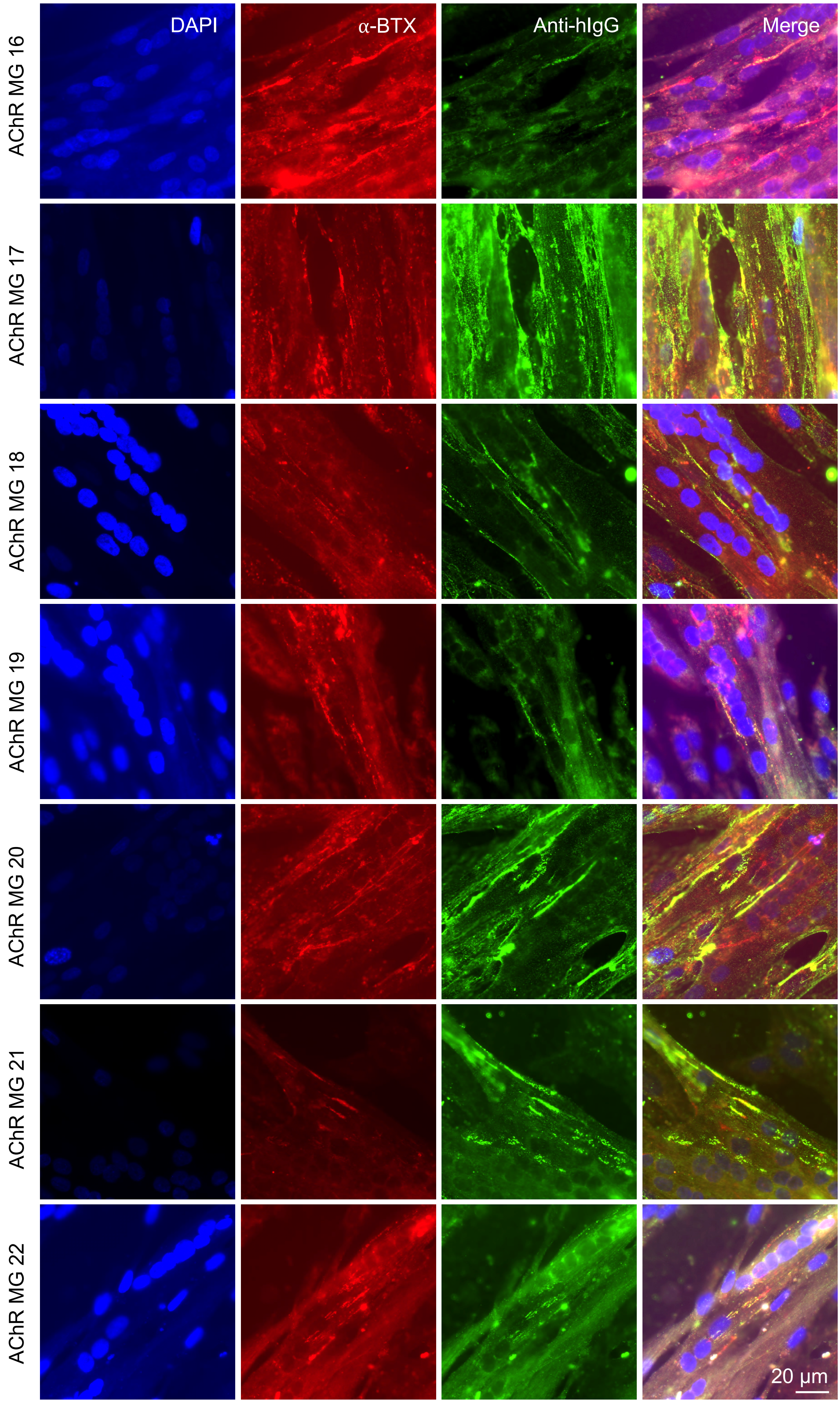



Continuation next page

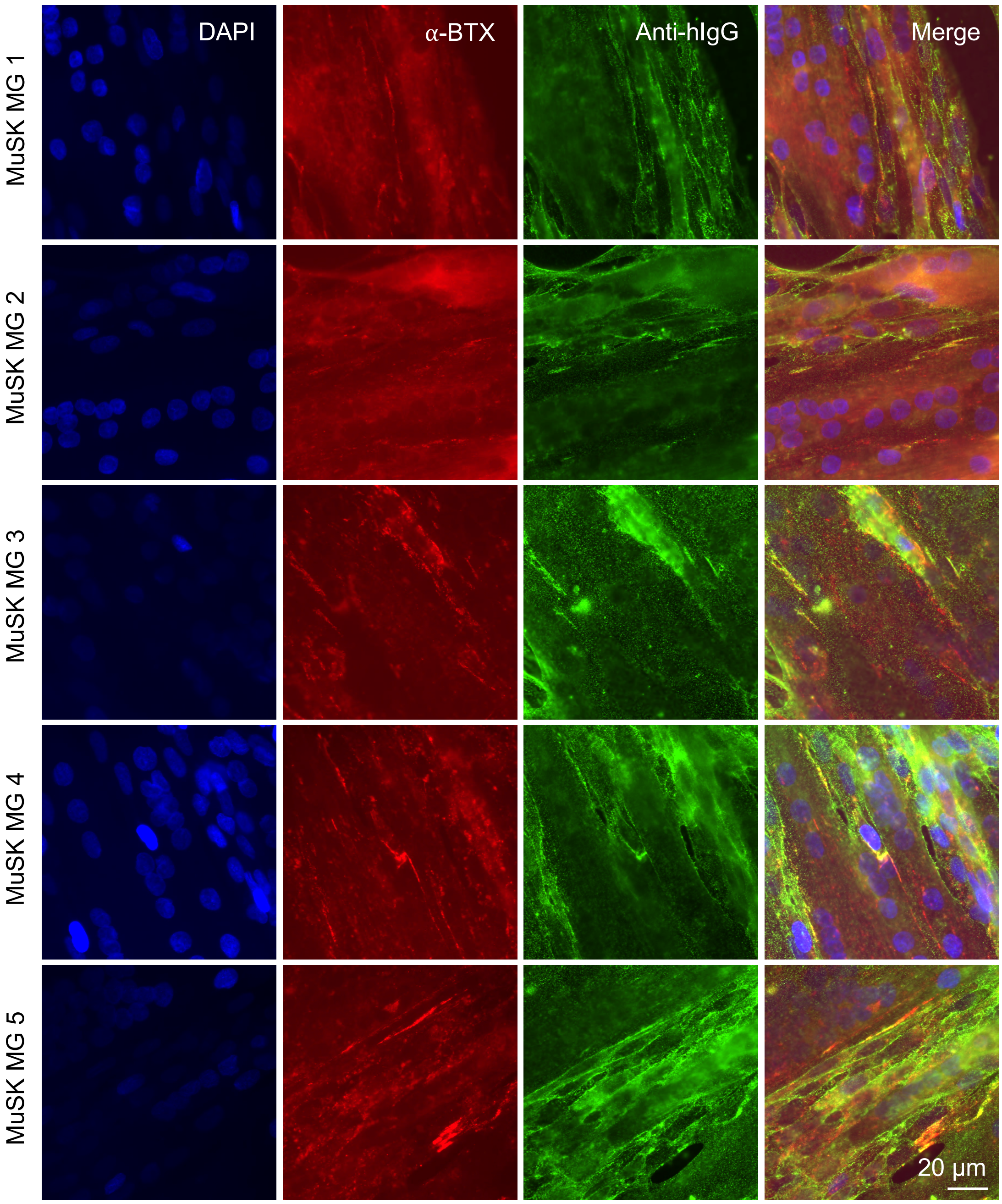


**Supplementary fig. S3: AChR- and MuSK-MG sera show antibody binding in proximity to AChR clusters.** Detection of colocalization of anti-human IgG (green channel, anti-human IgG-AF488) with the *in vitro* NMJs (red channel, α-bungarotoxin-AF594) by fluorescence microscopy. Blue staining = DAPI. Scale bar indicates 20 µm.

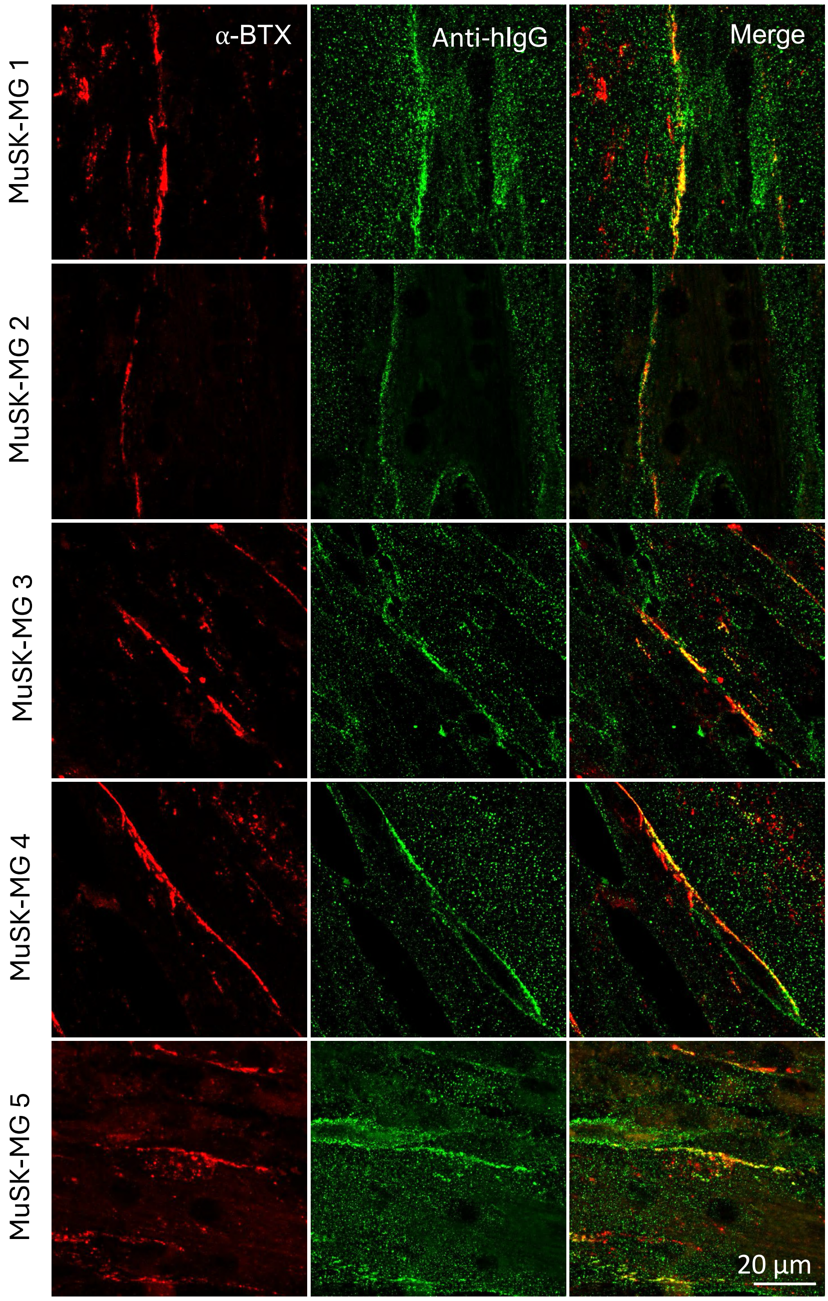


**Supplementary fig. S4: Validation of antibody binding to *in vitro* NMJs.** In patient samples with a weak signal in fluorescence microscopy (MuSK 1-6), antibody binding was validated by confocal microscopy, showing colocalization of anti-human IgG (green channel, anti-human IgG-AF488) and *in vitro* NMJs (red channel α-bungarotoxin-AF594). The z-stacks were flattened displaying the maximum intensity. Scale bar indicates 20 µm.
